# Fish from the sky: Airborne eDNA tracks aquatic life

**DOI:** 10.1101/2025.05.25.655195

**Authors:** Yin Cheong Aden Ip, Gledis Guri, Elizabeth Andruszkiewicz Allan, Ryan P. Kelly

## Abstract

Water and air are generally treated as separate reservoirs of environmental DNA (eDNA) derived from the species resident in those respective environments. However, it is likely that eDNA routinely crosses the air-water boundary in both directions as a result of deposition, evaporation, or other processes. Here, we systematically tested methods of sampling eDNA at the air-water interface, showing for the first time that aquatic life can be reliably detected from passive air samples collected nearby. We deployed four simple air samplers — three different kinds of filters and one open tray of deionized water — alongside paired water samples and visual counts over a six-week peak run of Coho salmon (*Oncorhynchus kisutch*) at a local spawning stream. We then quantified eDNA concentrations in both air and water (air: copies/day/cm2; water: copies/L) using quantitative PCR, to estimate (1) the concentration of target eDNA in air vs. water, and (2) the capture performance of each filter type. Despite an approximate 25,000-times dilution versus water, passive air collectors captured quantitative airborne eDNA signals that closely paralleled salmon counts, although recovery varied with sampler design and orientation. We show the air-water interface is a quantifiable source of aquatic genetic information using simple, passive samplers that do not require electricity, making them appealing for biomonitoring in remote or resource-limited settings. This work points the way to using airborne eDNA as a robust pathway for biological information critical to conservation, resource management, and public-health protection.

## 1 Introduction

Monitoring aquatic life is fundamental to understanding ecosystem health, guiding conservation efforts, and managing valuable natural resources Dudgeon et al. [2006], Reid et al. [2019]. Over the past decade, environmental DNA (eDNA) has upended our paradigm for biodiversity assessment by detecting and quantifying organisms through the genetic material they shed into their surroundings. Water-based eDNA surveys have proven particularly effective for monitoring endangered species (Biggs et al. [2015]), early detection of invasive species incursions (Thomas et al. [2020]), characterizing community composition (Wilkinson et al. [2024]), and even providing quantitative assessments for complementing traditional surveys (Allan et al. [2023], Guri et al. [2024b], Tillotson et al. [2018].

Meanwhile, airborne eDNA research has emerged as a promising frontier, though to date it has focused almost exclusively on terrestrial organisms. Both active and passive air filtration methods have been shown to recover DNA from mammals, birds, insects, and plants under field and enclosed conditions (Clare et al. [2021], Garrett et al. [2023], Johnson et al. [2019, 2023], Lynggaard et al. [2024], Roger et al. [2022], Lynggaard et al. [2022]). Intriguingly, these same air-sampling techniques sometimes detect strictly aquatic taxa. Yet in almost every case, these signals were never the focus of the study and are often treated as suspected contamination or attributed to zoo feed or piscivores fecal bioaerosols Sullivan et al. [2023], Klepke et al. [2022], Lynggaard et al. [2023, 2022], rather than recognized as a genuine ecological signal Tournayre et al. [2025].

Aquatic species signals detected in airborne eDNA surveys are often discarded Altermatt et al. [2025], Klepke et al. [2022], Johnson and Barnes [2024], Lynggaard et al. [2023]. Yet, these overlooked detections hint at a genuinely unexplored, rich, untapped source of data for a complementary biodiversity monitoring approach. Environmental systems are inherently interconnected Folke et al. [2021], and basic physics points to why aquatic DNA should appear in air Monahan et al. [1986], Seinfeld and Pandis [2016]. Natural physical processes at the air-water interface, such as evaporation, bubble-burst aerosolization from riffles and splashes, and leaping fish, churning substrates, create splash-driven aerosols; all of these provide plausible mechanisms for transferring eDNA into the atmosphere Duchemin et al. [2002], Mueller et al. [2008], Stell et al. [2020], Van Dijk et al. [2003]. These potential mechanisms would suggest that air could act as a diluted, but still informative, extension of the aquatic environment, representing biological signals that are real, quantifiable, and ecologically meaningful.

To test this hypothesis, we conducted the first targeted investigation of cross-medium water-to-air eDNA transfer, specifically examining whether genetic material from aquatic organisms can be detected in air samples collected above water surfaces. Leveraging the behavior of Coho salmon (*Oncorhynchus kisutch*) during their spawning season Mueller et al. [2008], we collected and measured eDNA concentrations in paired water and passive air samples over a six-week peak migration period, with visual fish counts from hatchery staff. To evaluate the mechanisms of airborne DNA capture and settlement, we deployed four passive air collection methods: vertically orientated gelatin air filters (commonly used in low-flow air filtration systems), polytetrafluoroethylene filters (PTFE; standard in high-flow air applications), and mixed cellulose ester filters (MCE; traditionally employed for water filtration), as well as an open, horizontal tray of deionized water exposed to ambient conditions Klepke et al. [2022]. These were chosen for their distinct physical properties, contrasting orientations, and different particle capture efficiencies targeting different aerosol types. By comparing detection sensitivity, temporal patterns, and quantitative performance of these four collectors against water eDNA concentrations and visual counts, we show that—even at roughly 25,000-times dilution—airborne eDNA can reliably track real-world aquatic population dynamics.

Our findings provide the first evidence that aquatic genetic material can indeed transfer from water to air under natural conditions, and extends further with quantitative demonstrations that reflect actual population dynamics with just passive air samplers. Far from weighing air against water sampling, this cross-medium detection and quantification represents not merely a technical advancement, but a conceptual leap in ecological monitoring: the previously ignored air–water interface now emerges as a measurable reservoir of aquatic genetic information. While this study centers on salmon, its implications extend broadly, proving that aquatic eDNA in the air is quantifiable and potentially species rich. This advancement re-imagines ecological monitoring as one that bridges water and air to build more resilient, comprehensive surveillance networks in a rapidly changing world. Such integrated approach can become particularly valuable during extreme environmental conditions–when droughts parch rivers, floods render wading unsafe, or public-health concerns close off contaminated waters Chen et al. [2024], Ruiz-Ramos et al. [2023], Wan et al. [2023]– indicating that airborne eDNA can offer a complementary pathway for capturing aquatic genetic signals.

## 2 Methods

### 2.1 Field Sampling

We conducted this study in Issaquah Creek, a salmon spawning stream, outside of the Issaquah Salmon Hatchery (47.529501° N, 122.039133° W) from 26 August 2024 to 18 November 2024, with nine sampling events. We chose nine field trips that spanned the entire coho salmon (*Oncorhynchus kisutch*) run, from the first arrivals in early fall of August, through peak abundance around 17 October 2024 and into the tail end of the migration in November. This schedule ensured that our eDNA sampling captured both the lowest and highest levels of fish activity. Within the same sampling period, visual fish counts were performed by the hatchery staff, and salmon escapement data were obtained from the Washington Department of Fish and Wildlife escapement reports (https://wdfw.wa.gov/fishing/management/hatcheries/escapement#2024-weekly). Throughout our nine sampling events, weather conditions ranged from clear, calm days to periods of rainfall; detailed records of air temperature, wind speed, relative humidity, and precipitation are provided in Supplementary Material 1.

Sampling for eDNA was conducted in 24-hour blocks with deployment and recovery around 9 a.m. (river water was collected only on the first day/timepoint). Four passive collection methods were evaluated: three filter types—gelatin (Sartorius, 47 mm diameter), PTFE (Whatman, 47 mm diameter), and MCE (Sterlitech, 5.0 *µ*m pore size, 45mm diameter)—and an open container of deionized water. The rationale for selecting these materials is as follows: gelatin filters are effective at capturing airborne particles and are particularly suited for applications where maintaining viability (e.g., for subsequent bacterial culturing; Wu et al. [2010]) is desired; PTFE filters are noted for their high durability and are widely used in active air sampling experiments Harnpicharnchai et al. [2023]; whereas MCE filters, typically employed for water filtration, were included to assess their performance in an airborne context Allan et al. [2023]. All three passive filters were deployed via custom 3D-printed “honeycomb” puck filter holders, based on open-source Thingiverse designs (IDs 4306478 and 979318; Zachary Gold, pers. comms.), and were suspended from a hatchery railing approximately 3 m above the river water level (Figure 1), with their collection surfaces oriented horizontally. All three passive filters were deployed via 3D-printed “honeycomb” puck holders, based on open-source Thingiverse designs (IDs 4306478 and 979318)^1^ and generously shared by Mautz et al. (2025).

**Figure 1:**
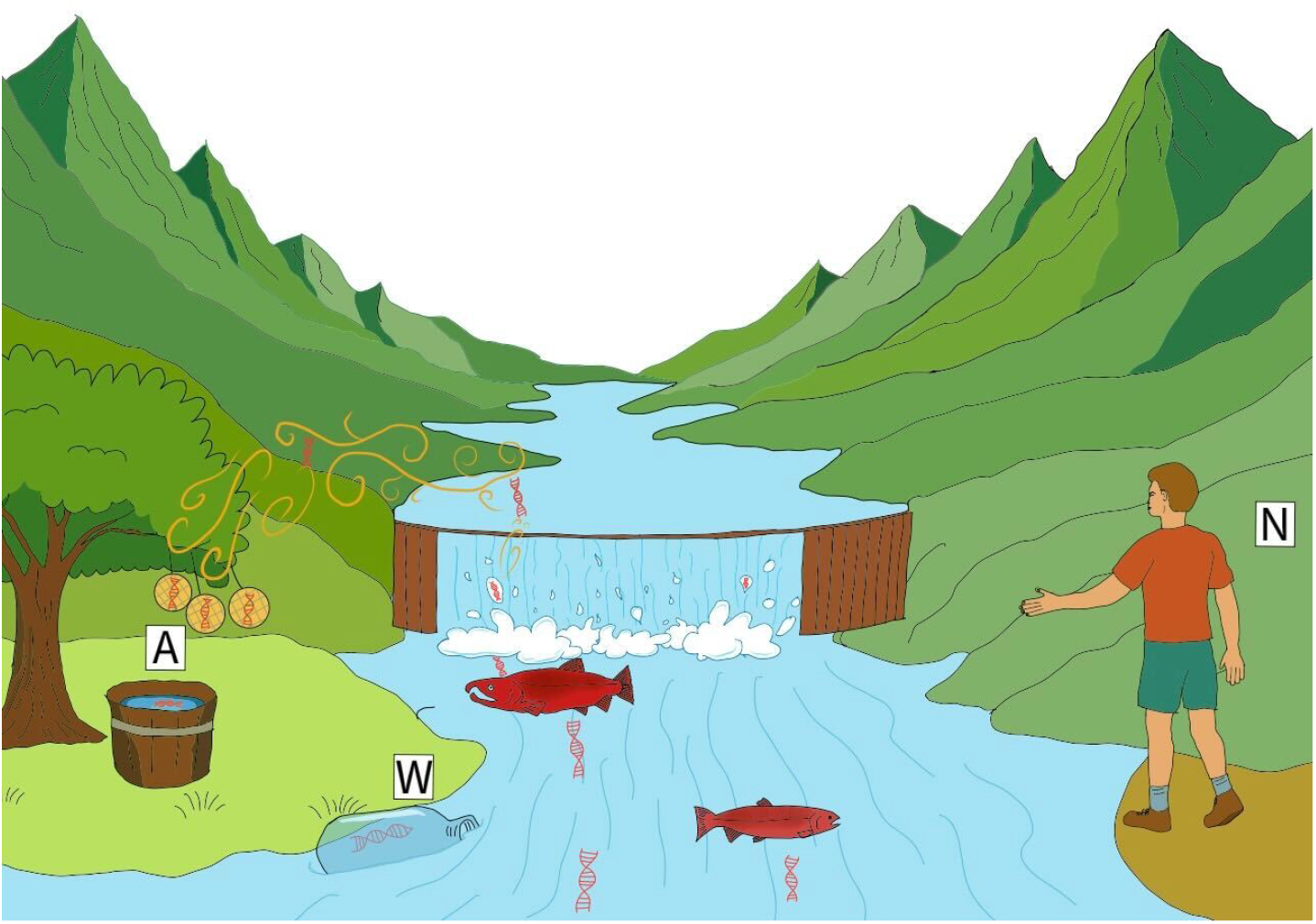
Conceptual illustration of cross-medium eDNA sampling above a salmon-spawning stream. Natural processes, including evaporation, bubble-burst aerosolization at riffles and splashes, and the vigorous movement of spawning Coho salmon, launch trace amounts of DNA from the river surface into the atmosphere. Passive airborne samplers, shown here as three vertically hung filters and an open tray of deionized water (A), intercept settling airborne eDNA, while paired water-grab sampling (W) and visual counts by hatchery staff (N) provide concurrent reference measurements.

This configuration minimizes splash contamination and captures airborne DNA particles driven by gravitational settling. The positioning relative to flowing water demonstrates how these passive collectors operate at a vantage point above the stream, facilitating non-invasive DNA capture in real-world field conditions. Two biological replicates were deployed for both gelatin and PTFE, while only one replicate was used for MCE. After the overnight deployment, filters were carefully recovered using sterile, disposable forceps and immediately immersed in 1.5 mL of DNA/RNA Shield.

The fourth passive collection method was an open container (25 cm width, 30 cm length, 10 cm depth) filled with 2 L of deionized water Klepke et al. [2022] also positioned about 3 m above the river and about 30 cm above the suspended passive filters. The open container was also deployed for about 24 hours at the same times as the passive filters. The container had an open surface (i.e., oriented horizontally) to capture airborne particles settling by gravity. The surface area of the container was approximately 750 cm^2^, in contrast to the roughly 16 cm^2^ surface area offered by the 47-mm circular filter disks. Only one replicate was used for the open water container method. Note that although 2 L of deionized water was placed in the container, heavy rainfall during overnight deployments sometimes increased the water level. Any debris (e.g., bugs, leaves) was removed from the container before on-site filtration using the same system as for river water samples (see below for details).

Concurrent with airborne sampling, water samples were collected by hand from the river directly adjacent to the coho salmon aggregation school, just downstream of the fish ladder entrance. The ladder is operated continuously by hatchery staff throughout the salmon run, ensuring fish passage and visual counts. A total of 3 L of water was filtered on site using a Smith-Root Citizen Science Sampler equipped with 5.0 µm mixed cellulose ester (MCE) filters Allan et al. [2023]. Three 1 L replicates were processed and immediately preserved in 1.5 mL of DNA/RNA Shield (Zymo Research) using clean disposable plastic forceps. Field-negative filtration blanks (1 L of Milli-Q water each) were also processed using the same system. All equipment was decontaminated with 10% bleach, thoroughly rinsed with deionized water, and handled with gloves to minimize contamination. All filters (passive air, passive open container, water) were stored at –20°C until processing, typically within one week.

### 2.2 Wet-Laboratory Procedures

DNA extraction from both river water samples and airborne filter samples was performed using the Qiagen Blood and Tissue Kit according to the manufacturer’s protocols. During extraction, it was noted that the gelatin filters dissolved completely in the DNA/RNA Shield. Samples were vortexed for 1 minute, and 500 *µ*L of the shield was used for DNA extraction. qPCR assays targeted a 114-bp fragment of the cytochrome b gene of coho salmon, using primers derived from Duda et al. [2021]— COCytb 980-1093 Forward: CCTTGGTGGCGGATATACTTATCTTA and COCytb 980-1093 Reverse: GAACTAGGAAGATGGCGAAGTAGATC. SYBR Green chemistry was employed using SYBR Select Master Mix (Fisher Scientific) on an Applied Biosystems QuantStudio 5 real-time PCR system with a 384-well block. Each 10 *µ*L reaction consisted of 5 *µ*L of SYBR Select Master Mix, 0.4*µ*L of 10 mM forward primer, 0.4 *µ*L of 10 mM reverse primer, 2.2 *µ*L of molecular grade water, and 2 *µ*L of DNA template. Melt curve analysis was performed to confirm the amplification of the target fragment, with an accepted melting temperature of 81°C ± 1°C.

Standard curves were constructed using a coho tissue DNA extract quantified with a Qubit fluorometer. The stock solution was diluted to 1.0 ng/*µ*L and designated as 10^6^ copies/*µ*L. Serial dilutions were then prepared, with 10^5^, 10^4^, and 10^3^ copies/*µ*L run in triplicate, 10^2^ copies/*µ*L in quadruplicate, and 10^1^/*µ*L copies in triplicate.

### 2.3 Joint statistical model

#### 2.3.1 Visual observation model

In summary, we synthesize three methods of observations (visual counting of fish, water eDNA measurements, and air eDNA measurements) which derive from a single unknown true fish accumulation rate (denoted here as *X*). Through this joint approach, we can estimate: the aerosolization factor (*η*; the magnitude of water eDNA transferred to air), the effectiveness and reliability of different passive air filtering techniques (*τ*), and the replicability of various filters (*ρ*) throughout the 6-week peak coho salmon spawning period.

We model the upstream migration of coho salmon (*Oncorhynchus kisutch*) as arising from the inferred unknown true density of fish. As individuals move upriver, fish accumulate in a holding area immediately downstream of the hatchery river dam. At discrete times *t*, the hatchery staff opens the ladder gate to allow passage into holding tanks. Between successive gate-opening events (Δ*t*), additional coho salmon arrive and join the backlog in the holding area, increasing the number of individuals awaiting passage. Let *X* represent the true daily accumulation rate (also fish density at the river dam) in units of fish/day at time *t*, and E denote the number of days elapsed between consecutive gate openings (*E*_*t*_ = Δ*t*; hence *E*_*t*=1_ = 0). Prior to each gate opening (at time *t*), the crew conducts a visual count *N*_*t*_ of accumulated fish. Assuming *X*_*t*_ remains relatively constant between successive gate-opening events (Δ*t*), we model the observed fish counts as a Negative Binomial process:

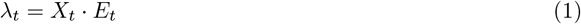

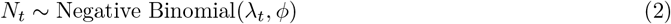

where, *N* is the visually observed number of fish at time *t, λ*_*t*_ is the expected number of fish accumulated over the *E*_*t*_-day interval with a fixed overdispersion parameter shared across time points (*ϕ* = 20; hence variance 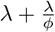) Welch and Ishida [1993], Guri et al. [2024b]

#### 2.3.2 Molecular process model

Let *W* be the unobserved eDNA concentration (copies/L) in the water at time *t* and let *ω* be the “integrated eDNA factor” – the conversion factor between *X* and *W* (see Guri et al. [2024b] for further interpretation of this parameter). We can express the relation between the fish density and water eDNA concentration as:

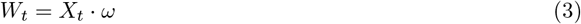

Let *A* be the unobserved eDNA concentration (copies/day/cm^2^) in the air at time *t* that is filtered using passive collection method *j*. We model the air eDNA concentration as a log-linear function of the water eDNA with intercept *η*, slope = 1, and error term *ε* (unexplained variability; time and filter specific):

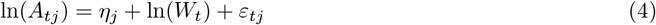

Here the intercept *η* can be interpreted as the water-to-air transferability (or dilution) factor and *ε* as the error term parameter (similar to the sum of squares error) from a linear regression where *ε*_*tj*_ ~ *𝒩* (0, *τ*_*j*_).

For some filter types *j* (PTFE filters (*j* = *P*); gelatin filters (*j* = *G*)) we sampled two biological replicates and we used the mean of those biological replicates to determine the average concentration in the air *A* at time *t* as following:

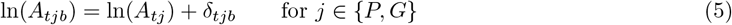

where *δ*_*b*_ indicates the deviation of individual biological replicate from the average concentration (*A*_*tj*_), following a normal distribution with mean 0, *δ*_*tjb*_ ~ *𝒩* (0, *ρ*_*j*_), with *ρ*_*j*_ indicating the magnitude of the replicates deviation from the mean sample. Because we have only two replicates at each sa led time, we impose a sum to zero constraint on the replicates (*b*) collected from a single sampling time (∑_*b*_ *δ*_*ij*_ = 0).

To estimate the levels of eDNA concentration in water (*W*) and air (*A*), we make use of the qPCR observation models (as described in Guri et al. [2024a], Shelton et al. [2022], with slight modifications). The model compartment uses the standard curve samples to estimate the intercept (*ϕ, β*0, *γ*0) and slope (*β*1, *γ*1) parameters between the known concentration (*K*) and the observed data (*Z* and *Y*) from qPCR machine as follows:

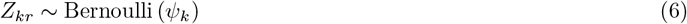

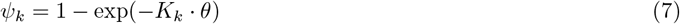

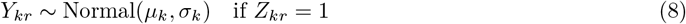

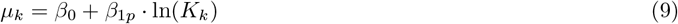

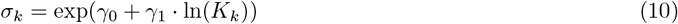

where *Z* is the binary outcome of target amplification for sample (*k*) and technical replicate (*r*) being present (1) or absent (0) following a Bernoulli distribution given the probability of detection *ψ* for each sample (*k*). The parameter *ϕ* is the intercept of the function between probability of detection *ψ* and the known DNA concentration (*K*; copies/*µ*L reaction) as the predictor variable. Additionally, for equations 8-10, *Y* is the observed cycle threshold (Ct) for sample (*k*) and technical replicate (*r*) which follows a normal distribution with mean *µ* (mean Ct) and standard deviation *σ* for each sample (*k*). We model *µ* as a linear function of known eDNA concentration (*K*) with intercept *β*0 and plate specific (*p*) slope *β*1 and the standard deviation *σ* of the observed Y as an exponential function of known eDNA concentration with intercept *γ*0 and slope *γ*1.

Subsequently, we build the same model compartment for estimating eDNA concentration in water and air by substituting *U*_*t*_ and *Q*_*tj*_ (and *Q*_*tjb*_ for *j* ∈ {*P, G*}) respectively, with *K*_*k*_ through equation 6-10 (see SI Appendix, Fig. S1), where *U* and *Q* are concentration normalized per reaction volume (*V* = 10*µ*L) and surface area (*S* = 16 cm^2^ for gelatin, PTFE, and MCE, and 750 cm^2^ for the open containers of deionized water) of *W* and *A* respectively as follows:

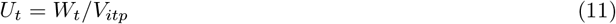

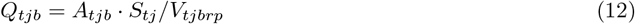

The intercept and slope parameters (from equation 7, 9, and 10) between qPCR observations and eDNA concentration of water, air and the standard samples are shared between model compartments.

### 2.4 Model conditions

The joint model (SI Appendix, Fig. S1) was implemented using the Stan language as implemented in R (package: Rstan) running four independent MCMC chains using 5000 warm-up and 5000 sampling iterations (for parameters and their prior distribution see SI Appendix, Table S1). The posterior predictions were diagnosed using statistics (Gelman and Rubin 1992) and considered convergence for values less than 1.05 and effective sample size (ESS) greater than 1000 for all parameters.

## 3 Results

Airborne eDNA passively detected Coho salmon and closely mirrored their abundance in the river during the upstream migration period. Detection efficiency and signal strength varied considerably among the different passive airborne eDNA capture methods.

### 3.1 Fish accumulation based on visual counts and eDNA in river water

Coho salmon migration occurs not as a single continuous event, but rather as a series of distinct burst peaks from mid-October through late November, where the peaks are highly likely to be connected with environmental factors such as water temperature and discharge. The average daily accumulation rate (X; black line in Figure 2) was estimated at 160.4 fish/day with peaks exceeding up to 286 fish/day and low activity of ca. 78 fish/day (Figure 2).

**Figure 2:**
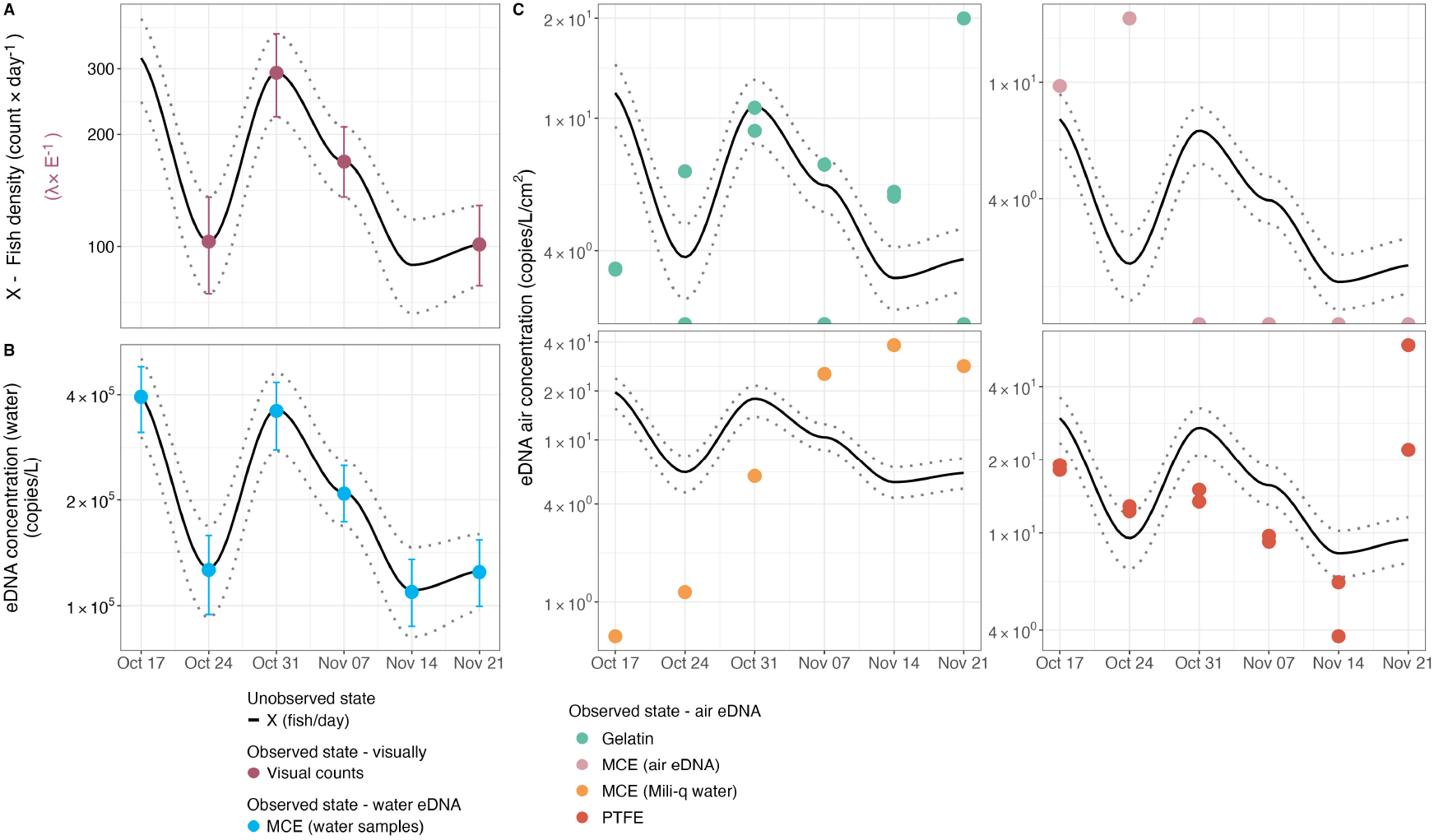
The temporal dynamics of estimated fish density in units of fish/day (X; black line with 95 confidence intervals - dotted lines) from October 17^*th*^ to November 21^*st*^ compared to the posterior distributions of visual observations (A), eDNA concentrations (copies/L) in water (B), and eDNA concentrations (copies/day/cm^2^) in air using various filter types (C).

Because both observation methods (river water eDNA and visual observation) are jointly used to estimate the daily accumulation rate (X), their concordance was best evaluated through the parameter *ω*. A converged and narrowly distributed *ω* parameter indicates strong agreement between the two methods and simultaneously a reliable conversion parameter from fish/day to eDNA copies/L. In this case, *ω* = 9.578 (95% quantile range of 9.352 to 9.804; (SI Appendix, Fig. S4)), suggesting consistent concordance between observed fish counts and eDNA concentrations hence, biologically, this implies that an accumulation rate of 1 fish/day corresponds to approximately 15000 (± 3000) copies/L.

### 3.2 Air eDNA signals

Airborne eDNA originating from coho salmon was successfully detected across all passive air collection methods deployed (gelatin, PTFE, MCE filters suspended in air, and open containers of deionized water - MCE DI water). These results provide compelling proof-of-concept evidence that genetic material from aquatic organisms can be recovered from the atmosphere without requiring active airflow systems, thereby demonstrating the viability of fully passive sampling approaches for detecting airborne aquatic eDNA under field conditions.

All filter types demonstrated that airborne eDNA concentrations were approximately 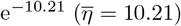 lower than corresponding waternborne eDNA concentrations, establishing a quantifiable dilution factor between water and air matrices. On average 1 copies/day/cm^2^ captured in air is equivalent to ca. 25,000 copies/L in water. Despite the general consistency in estimating the water-to-air dilution coefficient, method-specific variations in collection efficiency were observed (Table 1; SI Appendix, Fig. S2*A*). PTFE filters exhibited higher capture efficiencies, collecting 2 times more eDNA than the mean across all air sampling methods (Table 1). Deionized water tray (MCE DI water) demonstrated the second highest efficiency (1.3 times the average; Table 1), while gelatin filters and air suspended MCE filters showed comparatively lower capture efficacy (0.8 and 0.5 times the average, respectively; Table 1).

**Table 1:** Estimated posterior means of dilution parameter (*η*), standard deviation of the residuals (*τ*), biological replicability (*ρ*), and capturing efficiency 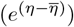.

| Filter type | Dilution<br>(in $\log_e$)<br>($\eta$) | Standard<br>deviation<br>($\tau$) | Biological rep.<br>error magnitude<br>($\rho$) | Capturing<br>efficiency<br>$e^{(\eta-\bar{\eta})}$ |
| --- | --- | --- | --- | --- |
| Gelatin | -10.45 | 0.473 | 0.386 | 0.785 |
| PTFE | -9.53 | 0.570 | 0.154 | 1.979 |
| MCE Air | -10.91 | 0.949 | - | 0.496 |
| MCE DI water | -9.95 | 1.780 | - | 1.295 |

In terms of alignment with the biological activity, PTFE and gelatin filters best mirrored the daily fish accumulation patterns, exhibiting the lowest residual error magnitude (expressed as the standard deviation of *ε*) with *τ* = 0.473 and 0.570, respectively (Table 1; SI Appendix, Fig. S2*B*). Conversely, MCE DI water, despite having the largest surface area, showed less agreement with the fish migration dynamics (*τ* = 1.780; Table 1; SI Appendix, Fig. S2*B*). The MCE air suspended filters performed least effectively in tracking temporal migration patterns, failing to amplify coho salmon DNA beyond the first two weeks of the sampling campaign Figure 2*C*.

Subsequently, biological replicates for gelatin and PTFE filters revealed additional insights regarding methodological robustness and reproducibility. PTFE filters produced the most consistent quantifications, with lower variance (expressed as the standard deviation of *δ*) between replicates (*ρ* = 0.154; Table 1; SI Appendix, Fig. S2*C*), whereas gelatin filters showed a higher degree of variability (*ρ* = 0.386; Table 1; SI Appendix, Fig. S2*C*), indicating reduced reproducibility of quantitative outcomes.

In sum, these performance differences across sampling methods likely reflect inherent physical and operational characteristics of each filter type and collection method, which in turn influence their ability to capture either discrete or cumulative biological signals from the source species.

### 3.3 Model diagnostics

Convergence and reliability of the Bayesian model were assessed through comprehensive diagnostics. All parameters (SI Appendix, Table S1) exhibited reliable Gelman-Rubin convergence statistics 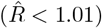 and effective sample sizes (ESS) exceeding 1000 per parameter, indicating successful convergence and efficient mixing of the six independent chains (SI Appendix, Fig. S3*A*).

No divergent transitions were detected during sampling, and the maximum tree depth was not exceeded, indicating no issues with divergence or exploration limits (SI Appendix, Fig. S3*B*). The posterior likelihood demonstrated convergence before the sampling phase began, with all chains exhibiting high mixing, confirming robust exploration of the parameter space (SI Appendix, Fig. S3*B*).

Prior sensitivity analyses revealed that posterior estimates differed from priors, demonstrating that the posteriors were appropriately updated based on the observed data rather than being heavily influenced by prior assumptions (SI Appendix, Fig. S4). Additionally, the posterior predictive checks (PPC) demonstrated that the model reliably reproduced the observed data, supporting the validity of parameter estimates (SI Appendix, Fig. S5).

Collectively, these diagnostics confirm the reliability and validity of the Bayesian model used here.

## 4 Discussion

### 4.1 A New Dimension of Biodiversity Monitoring

Our study establishes, for the first time, that passive airborne eDNA sampling can reliably capture and track molecular signals from aquatic organisms. This represents a paradigm shift in how we access aquatic biodiversity—through the air, without ever touching the water. In salmon-spawning streams, genetic material from the river is transported into the air likely by evaporation, bubble-burst aerosolization at riffles and splashes, and the rigorous churning of spawning fish Wood et al. [2021], Prather et al. [2013]. Collections from multiple passive samplers, when compared with conventional water-based eDNA assays and daily visual counts, reveal a clear and quantitative relationship between airborne eDNA concentration and salmon density.

Until now, airborne eDNA surveys have reported fish or other strictly aquatic taxa as likely to be laboratory contamination, zoo-feed artifacts, or piscivore fecal bioaerosols Klepke et al. [2022], Lynggaard et al. [2023], Sullivan et al. [2023], Lynggaard et al. [2022]. Our data suggest otherwise: this is a real ecological signal rather than an experimental artifact. It is likely that wind speed, relative humidity, and temperature will determine how far and how long aerosolized aquatic eDNA will travel before being redeposited Abrego et al. [2024], Giolai et al. [2024]. For example, high humidity and rainfall force rapid settling and very localized deposition while dry, windy conditions might carry genetic plumes further distances downwind Galbán et al. [2021], Maki et al. [2023]. These environmental controls, combined with air currents above riffles and splashes, should be considered when determining how passive samplers capture transient pulses of biological activity.

### 4.2 Sampler Design Shapes Signal Detection

Over 24-hour deployments, vertically oriented gelatin and PTFE filters acted as higher-resolution “fish-activity” samplers Jager et al. [2025]. Ambient air currents likely swept fine, splash-generated aerosols rich in salmon DNA onto their membranes, producing yields that rose and fell in sync with live fish counts and water-eDNA levels Blanchard and Woodcock [1980]. In contrast, the large horizontal tray of deionized water seemed to function more like a hydraulic-driven deposition trap. The tray saw a steady accumulation of eDNA over the six weeks and thus could have been collecting more coarse spray, foam, and decay-derived particulates from river turbulence and from decomposing carcasses Hinds and Zhu [2022], Prather et al. [2013]. Because salmon carcasses often remain in shallow banks and backwaters as the spawning season progresses, it is plausible that river turbulence and discharge could generate larger droplets over decomposing tissue, potentially facilitating eDNA dispersal Wood et al. [2021], Herman [2023]. These coarse droplets settle rapidly and dominate deposition on horizontal collectors while the fine fraction produced by active fish movement is under-represented. Gelatin and PTFE filters thus seem to provide snapshots of biologically driven eDNA flux whereas the water tray seemed to integrate both flow-driven and decay-driven inputs into a steadily rising accumulation curve.

The filter materials and operational context also shaped the air sampling performance. PTFE filters, known for their durability, delivered the most consistent results of all the passive filtration methods; gelatin filters yielded the highest sensitivity but showed greater variability; mixed cellulose ester filters captured negligible airborne DNA; and the open tray, with roughly 50 times more surface area than the vertical filters, recovered the highest total DNA yield. These samplers respond differently to weather. PTFE filters are generally unaffected by rain, whereas gelatin filters perform best in dry conditions because they dissolve when wet. Rainfall also introduces a trade-off for the tray: heavy rain can dilute the accumulated eDNA but may also scour additional airborne or splash-borne DNA into the water. Real-world debris, such as leaves, insects, sediment, can also wash into the tray, increasing the risk of clogging or necessitating pre-filtration before DNA extraction. Although our sample size is too small to quantify these opposing effects, future work should explicitly test how precipitation intensity influences both concentration and total yield Johnson et al. [2023].

### 4.3 Air is Dilute Water: Balancing Capture and Decay

On average, our data showed that eDNA in the air was roughly 25,000 times less concentrated than in the water, at a level that approaches the lower limits of qPCR detection. Despite this extreme dilution, our rigorous sampling, laboratory methods, and statistical models reliably picked up those sparse molecules, obtaining consistent, quantitative signals across the entire spawning season. To put this in perspective, dissolving a teaspoon of salt into a large aquarium yields a similar dilution magnitude: only a tiny fraction of shed DNA ever makes it into the overlying air, yet those few copies suffice to track real-time salmon activity. In particular, vertically oriented filters intercepted transient eDNA peaks that rose and fell in concert with visual counts, demonstrating that even at extreme dilution, airborne eDNA can still capture fine scale changes in fish presence that reflects real-world population dynamics.

Longer passive-sampler deployments naturally accumulate more settled eDNA, but they also expose collected material to ultraviolet radiation, microbial degradation and fluctuating humidity, all of which erode DNA integrity over time Brandão-Dias et al. [2023]. Strickler et al. [2015] showed that waterborne eDNA degrades with a half-life of hours to days under natural sunlight and microbial loads, suggesting that once deposited, airborne fragments may decay on comparable or even faster timescales. By contrast, Klepke et al. [2022] found that passive air particle collections continued to accrue new species detections for up to 96 hours, without specifying when deposition begins to be outpaced by degradation. In practical terms, this means that in a week-long deployment much of the DNA captured from early days may have degraded long before retrieval, so that the majority of signal reflects material deposited in the final one or two days of sampling. We therefore selected a 24-hour deployment to capture transient eDNA plumes while limiting post-deposition lossJohnson et al. [2023]. Future work should vary deployment lengths from a few hours to several days, and pair them with controlled decay assays, for example by spiking synthetic DNA onto filters and tracking its persistence, to determine the point at which accumulation and degradation balance.

Our findings also agree with the recent work by Jager et al. [2025] who showed that passive air samplers outperform active pumps by sampling intermittent, DNA-rich plumes over long intervals and detecting greater species richness. In our streams, vertical filters captured transient bursts of salmon eDNA in near real-time, while the open tray, and likely longer deployments, would tend to smooth those peaks into an integrated signal. Contrastingly, active-pump systems operating for only hours would average across plumes and risk overlooking fine-scale temporal changes.

By systematically mapping this interplay between deposition and degradation, and by benchmarking passive against active approaches, we can establish best-practice guidelines for airborne eDNA sampling durations. This effort would mirror how water-based eDNA workflows define optimal filtration volumes and storage times Barnes and Turner [2016], Altermatt et al. [2025] and would enable robust, context-specific deployments that maximize genuine signal recovery for real-time biodiversity monitoring.

### 4.4 Minimal Tools, Maximum Reach and Navigating Limitations

Perhaps most striking result from our work is the simplicity and versatility of passive airborne sampling. No pumps or power are needed, and equipment is extremely low cost and easy to deploy. This minimal-infrastructure approach makes it viable for remote headwaters, steep mountain channels, urban stormwater networks, and contaminated waters where sampling is unsafe Harrison et al. [2019], Bagley et al. [2019]. By integrating visual surveys, water-based eDNA and airborne eDNA, we establish a biomonitoring framework that is non-invasive, scalable and low-impact. In an era of intensifying droughts, floods and public-health risks such as bacterial outbreaks in stagnant waters, airborne eDNA offers resilient pathways for rapid invasive-species alerts, real-time disease surveillance in flood-prone wetlands and non-invasive population censuses in protected spawning grounds.

Naturally, challenges remain. Passive deployments rely on surface-area-by-time metrics rather than standardized air-volume units, complicating direct comparisons across studies. Optimal exposure times must balance accumulation against DNA degradation from UV, microbes and moisture Brandão-Dias et al. [2023]. Weather variability in wind, humidity and rain can alter deposition rates and sampler efficiency Johnson et al. [2023], Johnson and Barnes [2024]. Future work should refine sampler design, systematically compare vertical and horizontal orientations, explore automated or drone-based retrieval and integrate river discharge and meteorological data into quantitative models Galbán et al. [2021], Kirchgeorg et al. [2024], Shogren et al. [2017], Wood et al. [2021].

Our study begins to chart a portion of airborne eDNA ecology’s in five key dimensions Johnson and Barnes [2024]. We confirm origin by matching airborne DNA trends to co-occurring water eDNA and fish counts. We elucidate transport mechanisms such as evaporation, bubble bursts and fish activity. We quantify dispersal and dilution by measuring a 25,000-times concentration difference between water and air samples. We demonstrate fate through differential deposition on vertical filters and horizontal trays. Finally, we show that airborne DNA fragments remain amplifiable, offering an initial glimpse into their molecular state after transport. These insights lay the groundwork for future studies on persistence, degradation and particle-size distributions from airborne eDNA Brandao-Dias et al..

Overall, our work overturns the assumption that aquatic eDNA belongs solely underwater. By demonstrating that genetic signals from fish and other aquatic life routinely escape into and can be captured from the air, we open a new paradigm for ecological monitoring. The atmosphere above water emerges as a reliable, quantifiable reservoir of biodiversity data. This advance promises transformative applications from invasive species alerts in drought-stricken reservoirs to pathogen surveillance in flood-prone wetlands and non-invasive censuses in protected streams, equipping managers with a resilient, real-time window into aquatic ecosystems.

## Acknowledgments

We thank Natasha Kacoroski, Larry Franks, and the dedicated volunteers at Friends of the Issaquah Salmon Hatchery for their generous support in the field. We are also grateful to Travis A. Burnett and Darin Combs at the Washington Department of Fish and Wildlife for facilitating access and permitting field experimental work at the Issaquah Hatchery. Additional thanks to Pedro F.P. Brandão-Dias for assistance with air filter deployments, and to Kevan Yamanaka at the Monterey Bay Aquarium Research Institute for fabricating the 3D-printed passive filter holders. We are especially grateful to Chris Sergeant for valuable insights on salmon biology that shaped the interpretation of our findings, and to Ole Shelton for statistical advice that improved our analytical approach. We acknowledge funding support from the Packard Foundation [Grant No. GR016745].

## Author contributions

Y.C.A.I. conceived the study. Y.C.A.I. G.G and E.A.A. designed the field and laboratory protocols. Y.C.A.I. and G.G. jointly designed the downstream statistical analyses and Bayesian modeling framework. G.G. conducted all statistical analyses, with inputs from R.P.K. The fieldwork was performed by Y.C.A.I. and E.A.A., while Y.C.A.I. and G.G. co-wrote the manuscript. R.P.K. supervised the project, contributed to conceptual guidance, and provided critical revisions. All authors contributed to the study design and approved the final manuscript.

## Data availability

The authors declare that they have no competing interests. All data needed to evaluate the conclusions in this paper are available in the main text and/or the Supplementary Materials. Codes are available on https://github.com/gledguri/Air-eDNA-quant. Additional data, code, and materials will be made available upon reasonable request. No materials were subject to material transfer agreements (MTAs).

## Footnotes

1 https://www.thingiverse.com/thing:4306478; https://www.thingiverse.com/thing:979318

